# Deep proteome analysis of time-series human plasma samples applied to Covid-19 antibody test

**DOI:** 10.1101/2023.08.23.554537

**Authors:** Tetsuya Fukuda, Aya Nakayama, Yoko Chikaoka, Yasuhiko Bando, Takeshi Kawamura

## Abstract

For the purpose of Covid-19 antibody testing, the human plasma samples acquired over a period of 310 days from August 18, 2021, to June 22, 2022, were subjected to DIA LC-MS proteome analysis. This process led to the acquisition of a quantitative protein profile and allowed for the validation of the temporal behavior of expressed proteins within the samples. A total of 1502 proteins were identified from the plasma samples. Before vaccination, after the first dose, after the second dose, and during the period after contracting the novel coronavirus, protein quantification values during each event interval were compared. Despite minimal changes observed before and after Covid-19 vaccination, notable proteins exhibiting distinct high-expression and low-expression patterns were identified after contracting the novel coronavirus infection.

## Introduction

The novel coronavirus (Covid-19), discovered in 2019, demonstrated a formidable infectiousness and rapidly evolved into a global pandemic, spreading across the world and causing a high number of infections and fatalities. Researchers worldwide responded swiftly, focusing on the rapid development of treatments and vaccines. Approved medications were also quickly put into practical use and introduced to the market. However, similar to other viruses, Covid-19 has been generating new variants one after another. Despite these variants being attenuated in virulence, the situation remains uncertain and calls for continued vigilance [1-4].

Since the early stages, our research group has been conducting antibody tests for covid-19 within the human body and has acquired and stored a significant amount of human sample specimens. As a result, by subjecting plasma samples, particularly those taken before and after Covid-19 vaccination and infection, to in-depth proteomic analysis, aim to uncover the progression of pathogenesis through variations and behaviors of specific proteins indicative of immune responses, as well as protein fluctuation profiles that are likely to be identified.

In this study, blood sample collection began prior to the administration of the Covid-19 vaccine by a single research volunteer. Sequential sampling has been conducted, covering the first vaccine dose (Moderna), the second dose (Moderna), subsequent Covid-19 infection, and ongoing periodic sampling thereafter. Antibody testing has been carried out on these samples, along with the measurement and evaluation of antibody titers. Additionally, a time-series protein profile has been obtained through proteomic analysis.

For the proteomic analysis using LC-MS, applyed the DIA-LC-MS method [5-8]. Unlike conventional shotgun LC-MS proteomic analysis, where in the first stage only peptide-like spectra are selected from the MS spectrum of Single-MS and subjected to successive MSMS, this methodology has limitations in terms of the number of MSMS events due to the performance of the mass spectrometer. Weak peaks and similar components may be overlooked as a result. On the other hand, with DIA-LC-MS, in the DIA mode, for instance, by segmenting the MS range, such as m/z 425-445, into segments of 20 each, all the observed peaks in that range are collectively subjected to MSMS. As a result, there is a tendency for the identification count to increase compared to conventional shotgun LC-MS proteomic analysis.

We applied human plasma samples for Covid-19 antibody testing to the aforementioned analysis system, conducting LC-MS proteomic analysis. Through this process, obtained a quantitative protein profile and verified the behavior of expressed proteins within the sampled time series. In this report, we provide an account of our findings.

## Materials and Methods

### Samples

The samples used in this study consisted of 18 human plasma samples collected from a single volunteer donor through blood draws between August 18, 2021, and June 22, 2022. The management of these samples was conducted following institutional guidelines (The University of Tokyo) to ensure anonymity. The research involving these samples was approved by the Ethics Committee established at the University of Tokyo (Approval Number: 23-230). Details of the sample breakdown and antibody titers are provided in Table 1.

**Table 1.**
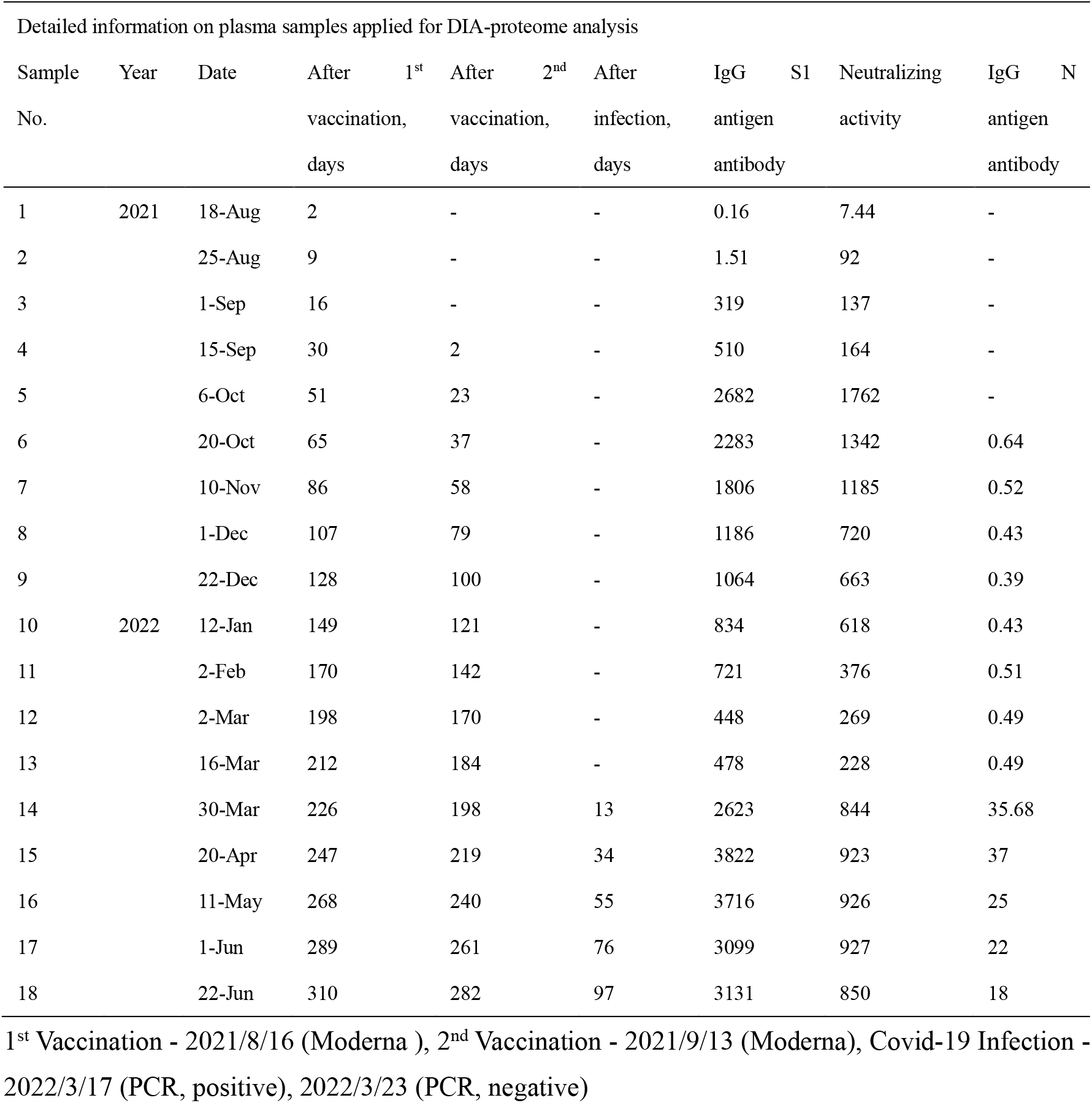
Information on the antibody titers and collection dates of the analyzed human plasma samples.

### Reagents

Protease inhibitor (Protease inhibitor Cocktail Tablets, complete, Mini, EDTA-free Tablets) were purchased from Roche Diagnostics (Indianapolis, USA). Benzonase (Benzonase® endonuclease, purity grade I (≥99%) suitable for biopharmaceutical production) was obtained from Merck (Darmstadt, Germany). Phosphate Buffered Saline (PBS), Ammonium bicarbonate (AMBIC), Dithiothreitol (DTT), Iodoacetamide (IAA), and Triethylammonium bicarbonate (TEAB) were acquired from Sigma (St. Louis, Missouri, USA). For the depletion of plasma samples, the Albumin and IgG Removal Kit (Amershan Biosciences, UK) was used. Sequencing-grade Trypsin was purchased from Promega (Madison, Wisconsin, USA). BCA reagent was obtained from ThermoFisher Scientific (Pierce Biotechnology, Madison, Wisconsin, USA). Acetonitrile, Trifluoroacetic acid, Formic Acid, and Methanol were procured from Kanto Chemical Co. Ltd. (Tokyo, Japan). SDS was purchased from Amersham Biosciences (Amersham, UK). Phosphoric Acid was obtained from NACALAI TESQUE, INC. (Kyoto, Japan). All water used in this study was produced using the Milli-Q system (Merck Millipore, Billerica, Massachusetts, USA).

For S-Trap processing, the following buffers and reaction solutions were prepared: 5% SDS in 50 mM TEAB (pH 7.55), 12% Phosphoric Acid, 90% Methanol in 100 mM TEAB (pH 7.1), 50 mM TEAB (pH 8), 0.2% Formic Acid and 0.2% Formic Acid in 50% ACN.

### Sample preparation

The major proteins in the plasma, such as Albumin and IgG, were removed using the Albumin and IgG Removal Kit (Amersham Biosciences, UK), applying the following procedure:

First, the resin in the vial was thoroughly mixed to create a suspension, and an appropriate amount (about 5 ml) was transferred to a 15 ml tube. Subsequently, the resin was washed three times with PBS. Adding 15 μL of plasma sample to 750 μL of the resin slurry, the mixture was shaken at room temperature for 30 minutes (1400 rpm, rt-21°C, 30 min → manual shaking at 250 rpm). The mixture was then subjected to centrifugation using a spin column (6500g, 5 min), and the flowthrough was collected. The BCA protein assay was conducted to quantify the protein amount, resulting in a total protein quantity of 2.1-2.3 mg.

Next, a TCA-Acetone precipitation was performed, and the collected precipitate was dissolved in 5% SDS in 50 mM TEAB (pH 7.55), bringing the total volume to 50 μL. Subsequently, the reduction and alkylation of the sample were carried out. 1M DTT was added to achieve a final concentration of 20 mM, followed by incubation at 95°C for 10 minutes and then returning to room temperature. Then, 1M IAA was added to achieve a final concentration of 40 mM, and the sample was incubated in darkness at room temperature for 30 minutes. After reduction and alkylation, 12% Phosphoric Acid was added to the sample solution to achieve a final concentration of 1.2%. 350 μL of 90% Methanol in 100 mM TEAB (pH 7.1) was added to bring the total volume to 405 μL. The sample was then applied to the S-Trap 96 well processing. The sample was gently transferred to the S-Trap 96 well plate, and 250 μL of 90% Methanol in 100 mM TEAB (pH 7.1) was added for protein retention and subsequent washing of the retained proteins. This operation was repeated four times consecutively. The Extrahera LV-200 (Biotage, Uppsala, Sweden) was used for these steps.

Then, 250 μL of 50 mM TEAB was added to 20 μg/vial of Trypsin (Promega) for dissolution, and immediately 125 μL of this solution was added to the S-Trap mini, which was then centrifuged at 4000g for 1 minute. After incubation at 47°C for 60 minutes, the reaction was continued overnight at 37°C. The S-Trap 96 well plate was set on the Extrahera LV-200, and 80 μL of 50 mM TEAB was added to recover the flowthrough. Subsequently, 80 μL of 0.2% HCOOH was added, and the flowthrough was collected. Furthermore, 80 μL of 0.2% HCOOH in 50% ACN was added, followed by centrifugation at 4000g for 1 minute to separate the flowthrough, which was then collected. After drying using a Speed Vac, the recovered samples were reconstituted in 200 μL of 0.1% TFA in 2% ACN.

### LC-MS conditions

The eluted samples were separated by nanoflow reverse-phase LC followed by analysis using a Orbi-Fusion mass spectrometer (Thermo Fisher Scientific, San Jose, CA) equipped with a Dream spray nano-electrospray ionization source (Dream spray, AMR Inc., Tokyo, Japan) [9-14]. The LC used was an Advance UHPLC (ultra-high-performance liquid chromatography) instrument (Bruker-Daltonics, Bremen, Germany) equipped with a HTS PAL auto-sampler (CTC Analytics AG, Zwingen, Switzerland). The samples were loaded onto a capillary reverse-phase separation column packed with 3.0-μm-diameter gel particles of pore size 120 Å (L-column2 ODS,3μm, 0.2x150 `mm`(CELI, Tokyo, Japan). Eluent A was 0.1% formic acid, and eluent B was 100% acetonitrile. The column was eluted at a flow rate of 1.2 μL/min with a concentration gradient was A + 5% B to 35% B in 100 min and from 35% B to 95% B in 1 min, with subsequent isocratic elution with 95% B for 8 min and a further concentration gradient from 95% B to 5% B in 1 min [15].

The mass spectrometer was operated in DIA (data-independent acquisition) mode in which the MS acquisition with a mass range of m/z 495-745 was automatically switched to MS/MS acquisition under the automated control of X calibur software 3.1 (Thermo Fisher Scientific).

DIA were collected using staggered windows with a loop count of 20.0 m/z isolation windows and collision energy mode stepped with 5 ms maximum injection time at custom mode.

In the DIA mode, each cycle consisted of a MS1 scan of 495–745 m/z with 60,000 resolution and AGC target of 2 × 106, followed by 50 MS2 scans of 498–742 m/z with resolution of 30,000 and AGC target of 2.5 × 10^6^ [7,8,16].

### DIA Data Processing

Acquired raw data were processed by DIA-NN (Data-Independent Acquisition by Neural Networks) for proteomics analysis. DIA-NN is known to be particularly useful for high-throughput proteomics applications because it improves the performance of protein identification and quantification in traditional DIA mode proteomics applications, enabling fast and reliable protein identification [6,17].

The algorithm for DIA-NN is described as follows. DIA-NN version 1.8.1, library free mode was used with the same Uniprot FASTA database. The Protein Database was applied to the Human database (UniProt Reference Proteome - Homo sapiens, Taxonomy 9606 - Proteome ID UP000005640 -. 20373 entries - UniProt release 2022_03, reviewd human cannonical).

Precursors of charge state 1–4, peptide lengths 7–30 and peptide m/z 300–1800 were considered with maximum one missed cleavage. A maximum of one variable modification per peptide was considered. Cysteine carbamidomethylation enabled as a fixed modification, N-terminal methionine excision as a variable modification, methionine oxidation as a variable modification, and N-terminal acetylation as a variable modification. Precursor FDR were then filtered at 1% False Discovery Rate [6,7,8,16,17]. SimpliFi™ (Protifi, LLC, USA) was used for quantitative analysis and evaluation of protein profiles.

## Results and discusstions

Using the aforementioned methods, sample preparation and LC-MS-based DIA proteomic analysis were conducted to acquire the protein profile of plasma samples. A total of 1502 human proteins were identified from the 18 plasma samples. When comparing the results of DIA-LC-MS proteome analysis between before the first vaccination (Moderna) and the sample after Covid-19 (variant) infection, particularly in sample number 1 and 2, interesting protein variations were observed, as validated using the SimpliFi™ software. Among these proteins, 60S Ribosomal protein L11 (RL11_human, p62913) stood out (Fig. 2).

**Fig. 1.**
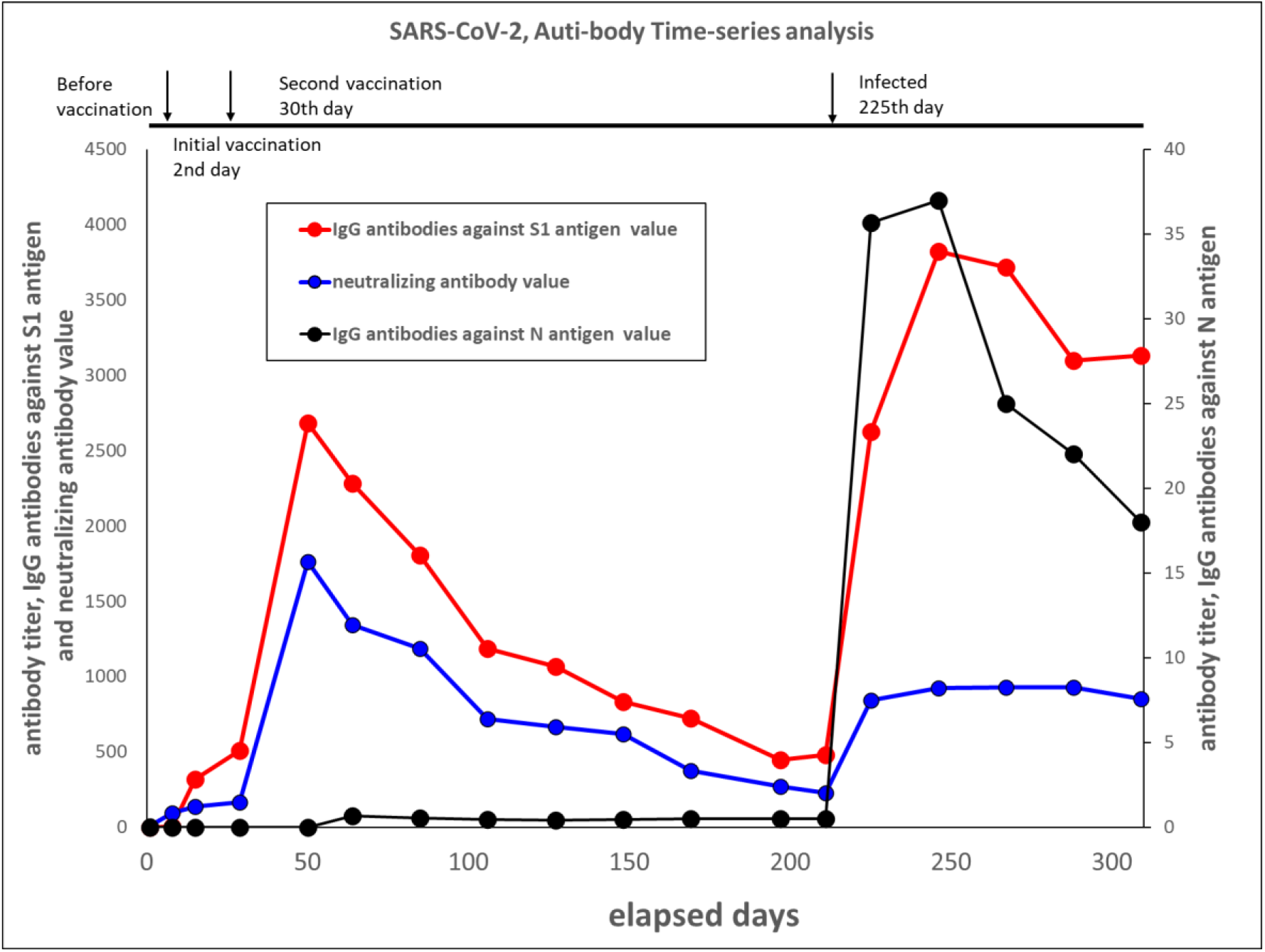
Fluctuations in the antibody titers against SARS-CoV-2 in human plasma samples. Antibody titers in human plasma samples from a single donor were measured over a period of 310 days, from August 18, 2021, to June 22, 2022. The donor received their first dose of the vaccine (Moderna) two days after the initial blood collection, followed by the second dose 30 days later (Moderna). After 225 days from the sample collection date, the donor became infected with Covid-19.

**Fig. 2.**
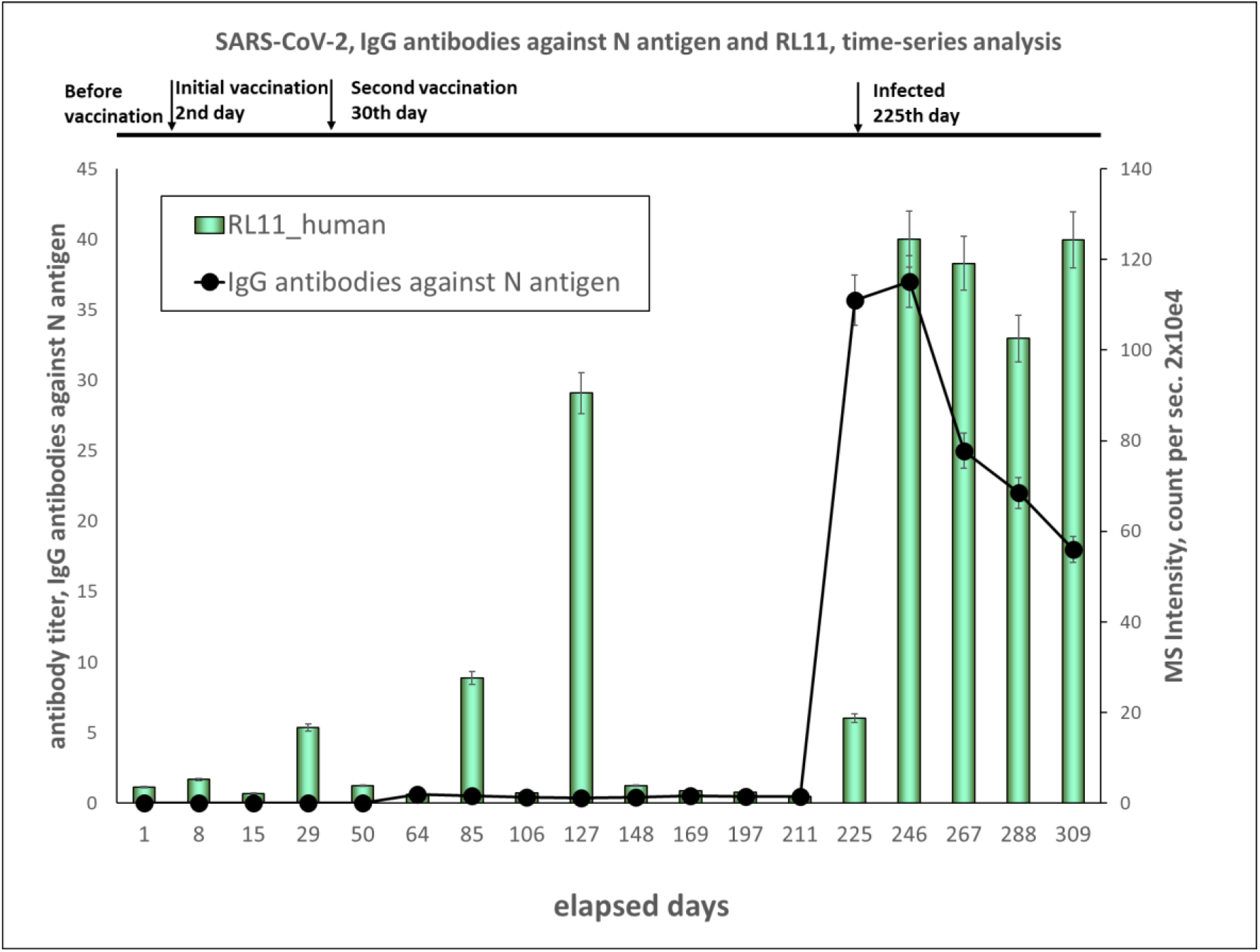
Temporal Changes in Quantitative Levels of IgG Antibodies against N Antigen and RL11_human in Human Plasma Samples This graph was plotted the quantitative values from proteome analysis of IgG antibodies against N antigen (which do not increase without Covid-19 infection) and RL11_human. Following Covid-19 infection, observed a significant upregulation of RL11_human.

While RL11_human did not exhibit significant changes after 1st and 2nd vaccination, it displayed a clear phenomenon of upregulation following Covid-19 infection, correlating with changes in IgG N antigen antibody levels. Investigation into this phenomenon revealed previous reports of a similar significant increase in the expression of this 60S Ribosomal protein after Covid-19 infection. Z. Khalid et al. have previously reported that 60S Ribosomal protein L29 (RPL29) was highly expressed across all COVID-19 infection groups [18-20]. The screened 60S ribosomal protein L11 (RL11_human) in this analysis is a component of the 60S ribosomal subunit, a central part of the ribosome, with several important functions: Ribosome conformation: RL11_human plays a crucial role in the conformation and maturation of ribosomes. It has a central role in the formation of the 60S ribosomal subunit, interacting with other ribosomal proteins to contribute to the accurate construction and structural formation of the ribosome [21]. RL11_human is also involved in the regulation of transcription control [22]. Under certain circumstances where ribosome synthesis decreases, immature ribosomal subunits are formed. In this state, RL11_human binds to transcription factors in the nucleus, regulating transcription. This mechanism is crucial for balancing ribosome synthesis and mRNA in the cell [23]. Regulatory Role: RL11_human acts as a regulatory factor in ribosome formation and function. For example, it interacts with p53 and is involved in p53 activation and cellular stress response [24,25]. RL11_human also is related in the control of apoptosis (cell death) [25].

The identification of this RL11_human in this analysis, along with its alignment with the variation profile of the antigen antibody group from the antibody testing, suggests relevance to previous reports [18-20]. According to the report by Q. Zhu et al. [19], they propose the potential to create a conducive cellular environment that significantly facilitates cell replication and promotes a favorable context for virus replication. They have elucidated that the expression of 60S Ribosomal protein RPL18 notably increases following Covid-19 infection. This finding suggests that the heightened expression of RPL18 is associated with cellular proliferation. Although RPL18 wasn’t directly observed in the dataset of this study, the similarity between the high expression of a related protein, RL11_human, and the current findings implies a possible connection. This suggests that the increased expression of RL11_human might share a similar role related to cell proliferation and virus replication.

On the other hand, U11/U12 small nuclear ribonucleoprotein 35 kDa protein (Q16560, U1SBP_HUMAN) were identified to exhibit a decline in detection levels following vaccination and were not detected after infection (Fig. 3). This U1SBP_HUMAN is a protein involved in regulating gene expression by controlling transcription and mRNA processing of the human genome. It functions as a nuclear ribonucleoprotein and is a crucial factor contributing to the accuracy and regulation of gene expression[26-29]. Aberrant dynamics or mutations of this protein suggests that associated with disorders related to erroneous splicing and abnormal gene expression. In this study, post-infection, low expression and potential functional impairment within the body were suggested; however, further verification is necessary to elucidate the specific impact. In the future, we will further scrutinize other molecules and strive to obtain detailed data regarding the variations in protein profiles due to SARS-CoV-2 infection.

**Fig. 3.**
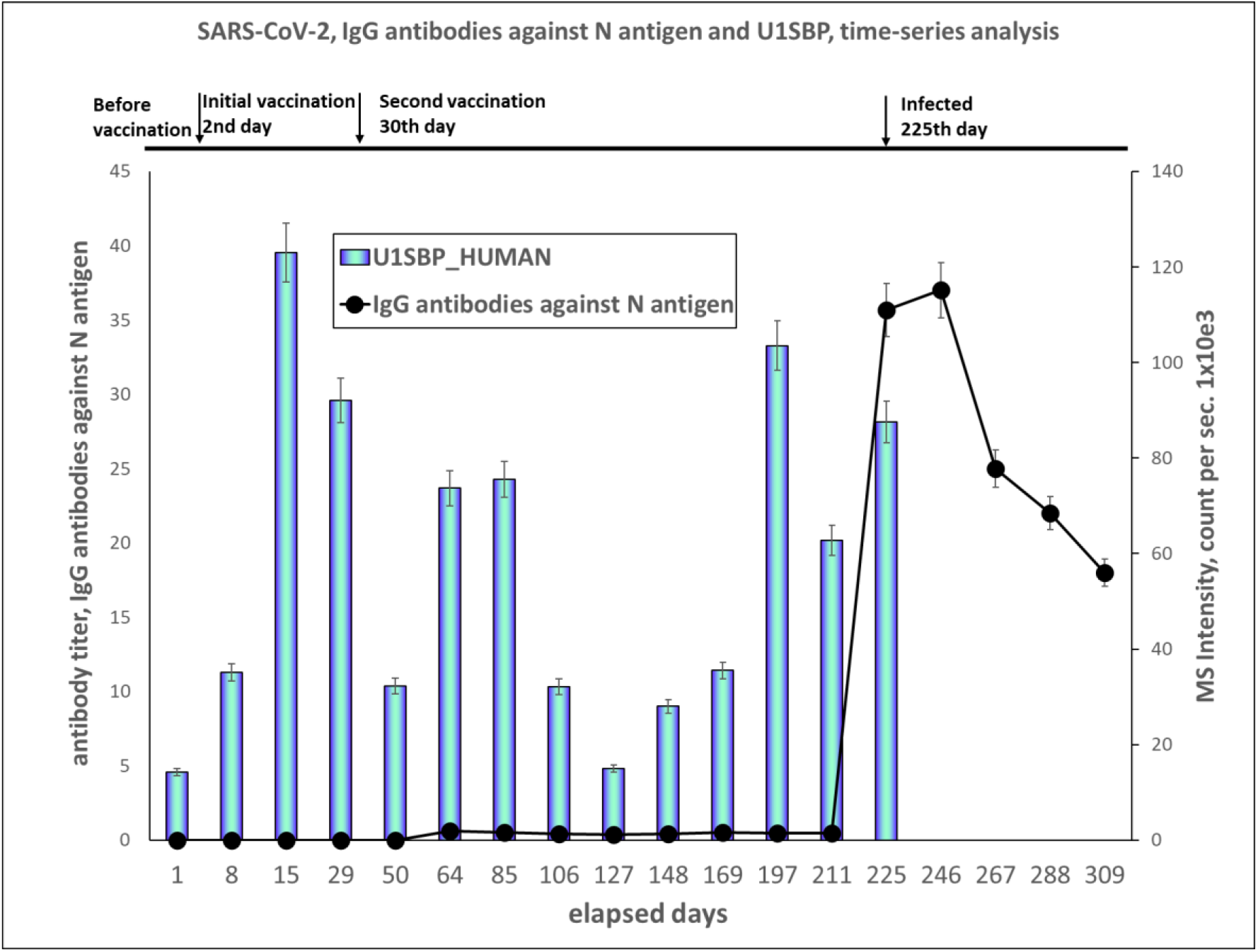
Temporal Changes in Quantitative Levels of IgG Antibodies against N Antigen and U1SBP_human in Human Plasma Samples This graph was plotted the quantitative values from proteome analysis of IgG antibodies against N antigen (which do not increase without Covid-19 infection) and U1SBP_human. Following Covid-19 infection, observed that U1SBP_human decreased to below the detection limit in the proteome analysis.

